# Distinct neurophysiological correlates of the fMRI BOLD signal in the hippocampus and neocortex

**DOI:** 10.1101/2021.02.01.429258

**Authors:** Paul F. Hill, Sarah E. Seger, Hye Bin Yoo, Danielle R. King, Bradley C. Lega, Michael D. Rugg

## Abstract

Functional magnetic resonance imaging (fMRI) is among the foremost methods for mapping human brain function but provides only an indirect measure of underlying neural activity. Recent findings suggest that the neurophysiological correlates of the fMRI blood-oxygen-level-dependent (BOLD) signal might be regionally specific. We examined the neurophysiological correlates of the fMRI BOLD signal in the hippocampus and neocortex, where differences in neural architecture might result in a different relationship between the respective signals. Fifteen human neurosurgical patients (10 female, 5 male) implanted with depth electrodes performed a verbal free recall task while electrophysiological activity was recorded simultaneously from hippocampal and neocortical sites. The same patients subsequently performed a similar version of the task during a later fMRI session. Subsequent memory effects (SMEs) were computed for both imaging modalities as patterns of encoding-related brain activity predictive of later free recall. Linear mixed-effects modelling revealed that the relationship between BOLD and gamma-band SMEs was moderated by the lobar location of the recording site. BOLD and high gamma (70-150 Hz) SMEs positively covaried across much of the neocortex. This relationship was reversed in the hippocampus, where a negative correlation between BOLD and high gamma SMEs was evident. We also observed a negative relationship between BOLD and low gamma (30-70 Hz) SMEs in the medial temporal lobe more broadly. These results suggest that the neurophysiological correlates of the BOLD signal in the hippocampus differ from those observed in the neocortex.

**Significance Statement:** The blood-oxygen-level-dependent (BOLD) signal forms the basis of fMRI but provides only an indirect measure of neural activity. Task-related modulation of BOLD signals are typically equated with changes in gamma-band activity; however, relevant empirical evidence comes largely from the neocortex. We examined neurophysiological correlates of the BOLD signal in the hippocampus, where the differing neural architecture might result in a different relationship between the respective signals. We identified a positive relationship between encoding-related changes in BOLD and gamma-band activity in frontal, temporal, and parietal cortex. This effect was reversed in the hippocampus, where BOLD and gamma-band effects negatively covaried. These results suggest regional variability in the transfer function between neural activity and the BOLD signal in the hippocampus and neocortex.

## Introduction

Functional magnetic resonance imaging (fMRI) is one of the foremost noninvasive methods for the examination of human brain function. However, despite the near-ubiquity of fMRI in cognitive neuroscience research, the blood oxygen level dependent (BOLD) signal, the basis of fMRI, provides only an indirect measure of underlying neural activity. Prior studies that acquired simultaneous fMRI BOLD and intracranial electrophysiological (iEEG) recordings from primary sensory cortices of non-human mammals have consistently reported that stimulus elicited BOLD signal changes are strongly correlated with changes in high frequency (> 30 Hz) gamma-band activity measured in extracellular local field potentials (LFPs) (Goense & Logothetis, 2008; Logothetis et al., 2001; Niessing et al., 2005). Subsequent multimodal imaging investigations in humans have largely confirmed the close relationship between changes in BOLD signal intensity and high frequency LFPs in auditory (Nir et al., 2007), sensorimotor (Hermes et al., 2012), and association (Conner et al., 2011; Ojemann et al., 2010) cortices.

The relationship between the fMRI BOLD signal and its underlying neurophysiology has generally been assumed to be uniform across different brain regions. Recent findings challenge this assumption, however, raising questions about the possible regional specificity of coupling between BOLD and LFP signal modulations (Conner et al., 2011; Ekstrom et al., 2009; for reviews, see Ekstrom, 2010, 2020; Logothetis, 2008; Ojemann et al., 2013). Of particular relevance to the current study is the potential for a dissociation between the fMRI BOLD signal and the underlying neurophysiology in the hippocampus, where sparse vascularization and neural coding schemes might lead to a different relationship between the respective signals evident in the neocortex (for review, see Ekstrom, 2021). This possibility is strengthened by the very different laminar organizations that are found in hippocampal allocortex and the neocortex, including neocortical regions adjacent to the hippocampus such as the entorhinal and parahippocampal cortices.

In the only multimodal fMRI-iEEG study of the human medial temporal lobe (MTL) to date, Ekstrom and colleagues (2009) compared measures of fMRI BOLD signal with extracellular iEEG activity recorded from the hippocampus and parahippocampal gyrus in five neurosurgical patients as they performed a virtual navigation task. A positive correlation between changes in BOLD signal and theta (4-8 Hz) activity was evident in the parahippocampal gyrus and, to a weaker extent, the hippocampus proper. Crucially, and in contradiction to the aforementioned findings from sensory and association cortex, changes in high frequency gamma activity did not correlate significantly with corresponding BOLD activity in either the hippocampus or parahippocampal gyrus. It bears mentioning however that these findings were based on a small sample of subjects (n = 5) with recordings confined to the MTL. It is therefore unclear whether the lack of correlation between BOLD and high frequency LFPs was the result of insufficient power, and whether potential BOLD-LFP coupling in the hippocampus and proximal MTL structures truly differed from that observed on the cortical surface.

In the present study, 15 patients with medically resistant temporal lobe epilepsy (TLE) implanted with depth electrodes performed a verbal delayed free recall task while iEEG was recorded simultaneously from hippocampal and neocortical sites. The same patients subsequently performed a similar version of the free recall task in a later fMRI session (Hill et al., 2020). Subsequent memory effects (SMEs) were computed for fMRI and iEEG as patterns of encoding-related brain activity that were predictive of successful recall following a brief distractor interval (Paller & Wagner, 2002). fMRI BOLD SMEs extracted from distributed hippocampal and neocortical sites were correlated with electrophysiological SMEs obtained from the same sites. The primary aim of the study was to identify the iEEG frequency band(s) that best predicted a commensurate BOLD response, and to determine whether the relationships between BOLD and iEEG SMEs varied between the hippocampus and neocortex.

## Materials and Methods

Behavioral and group-level fMRI data from this experiment were the topic of a prior report (Hill et al., 2020). The present descriptions of the free recall task and behavioral results overlap heavily with the descriptions given in that report and are only summarized here. The fMRI and iEEG findings described below have not been reported previously.

### Participants

Fifteen patients with medically resistant temporal lobe epilepsy were recruited to participate in this experiment (21-59 years, *M* = 37 years, *SD* = 12 years, 10 females). Three participants were left-handed, and all spoke fluent English before the age of five. Each patient underwent iEEG to localize and monitor epileptogenic activity, during which time they performed a verbal delayed free recall task similar to the one performed during a subsequent fMRI session. The number and placement of the electrodes were determined solely on the basis of clinical considerations. Origin of epileptogenic activity was right lateralized in seven patients, left lateralized in four patients, and bilateral in the remaining four patients. Enrollment was limited to patients who correctly recalled at least 10% of study items across a full iEEG session. No patient had radiological evidence of hippocampal sclerosis. The average delay between iEEG surgery and the fMRI session was 87 days (SD = 66 days). All patients gave informed consent in accordance with the University of Texas at Dallas and University of Texas Southwestern Institutional Review Boards and were financially compensated for their time.

### Free Recall Task

Patients performed similar versions of a verbal delayed free recall task while undergoing iEEG recording and fMRI scanning on separate occasions. All patients completed the iEEG version of the experiment prior to enrolling in the fMRI study. Both versions of the recall task comprised three phases: study, arithmetic distractor, and free recall (see below for session specific parameters). During the study phase, participants viewed words randomly selected from a database of high frequency concrete nouns (https://memory.psych.upenn.edu/WordPools). All words were concrete nouns between three and six letters in length, with a mean frequency per million of 46.89 (SD = 84.37, range 0.55 to 557.12) obtained from the SUBTLEX-US corpus (Brysbaert & New, 2009). Concreteness ratings ranged between 3.75 and 5 (*M* = 4.80, *SD* = .20) on a scale from 1 (most abstract) to 5 (most concrete) (Brysbaert et al., 2014). Participants were instructed to form a mental image of the object denoted by each word and to refrain from saying the word aloud or rehearsing previously studied words. The study phase was followed by a brief arithmetic distractor task to prevent rehearsal and to clear the contents of working memory.

Immediately following the distractor interval, participants were prompted to freely recall as many words from the immediately preceding study list as they could remember, in any order, for 30 seconds. Responses were made verbally and transcribed for subsequent analyses.

### fMRI Session

Participants received instructions on the experimental tasks and performed several practice trials prior to entering the scanner. During the task proper, they completed a total of 18 Study-Distractor-Recall cycles divided equally over six functional scanner runs. Structural T1 MPRAGE scans were collected upon completion of the final block. The entire scanning session took approximately 65 minutes. During the study phase, participants viewed lists of 15 words presented sequentially in white font on a black background. The presentation of each word was preceded by a red warning fixation cross presented for 500 ms, followed by the presentation of a single word for 1800 ms. An additional seven null trials (white fixation cross) were pseudo-randomly interspersed throughout each study list under the constraint that no more than three null trials occurred consecutively. This resulted in an inter-stimulus fixation interval that jittered between 900 and 9600 ms. Immediately following the study phase participants performed a 15s distractor task involving simple arithmetic problems in the form of ‘A+B=C?’. Participants were tasked with indicating whether the expression was correct or incorrect via a button press using their right index and middle fingers (counterbalanced across participants). Each expression remained on the screen until a response was made, with the instruction that responses should be made quickly and accurately. Verbal responses during the free recall phase were recorded for later transcription using a scanner-compatible microphone (Optoacoustics) and noise-cancelling software (OptiMRI v. 3.2) to filter out scanner noise.

### iEEG Session

All patients performed a version of the free recall task similar to that described above for the MRI session with the following differences. Patients performed 26 Study-Distractor-Recall cycles per session (the first of these being for practice and not included in the analyses). Seven of 15 patients completed more than one session (Mean # sessions = 3, range = 2-7), with multiple sessions per patient occurring on average two days apart. The task was performed on a laptop computer during an inpatient hospital stay following intracranial electrode placement. Study lists were composed of 12 concrete nouns selected at random without replacement. Four patients completed a protocol that included 10 items per study list; for these subjects the data analyzed came from an experiment that included brain stimulation, but only lists in which all items were presented and recalled in the absence of stimulation (non-stimulation lists) were included in the analyses. Each word was presented for 1800 ms followed by a random inter-item fixation jitter (750-1000 ms). Following each study list, patients performed a 20 s arithmetic distractor task comprising expressions in the form of ‘A+B+C+?’. Patients were required to enter a response to each expression via the keyboard. The free recall phase was identical to that described for the MRI session.

### MRI data acquisition and preprocessing

Functional and anatomical images were acquired with a 3T Philips Achieva MRI scanner (Philips Medical Systems, Andover, MA, USA) equipped with a 32-channel receiver head coil. Functional images were acquired using a T2*-weighted, blood-oxygen level-dependent echoplanar (EPI) sequence (sensitivity encoding [SENSE) factor 2, flip angle 70 deg, 80 x 78 matrix, field of view [FOV) = 24 cm, repetition time [TR) = 2000 ms, and echo time [TE) = 30 ms). EPI volumes consisted of 34 slices (1-mm interslice gap) with a voxel size of 3×3×3 mm. Slices were acquired in ascending order oriented parallel to the anterior commissure-posterior commissure line. Each functional run included 201 EPI volumes. T1-weighted anatomical images were acquired with a magnetization-prepared rapid gradient echo pulse sequence (FOV = 240 x 240, 1 x 1 x 1 mm isotropic voxels, 34 slices, sagittal acquisition). Participants performed a total of 18 study-test cycles split evenly into six scanner runs.

All fMRI preprocessing and analyses were conducted with Statistical Parametric Mapping (SPM12, Wellcome Department of Cognitive Neurology, London, UK), run under Matlab R2017a (MathWorks). Functional images were realigned to the mean EPI image and slice-time corrected using sinc interpolation to the 17^th^ slice. The images were then reoriented and spatially smoothed with an isotropic 8 mm full-width half maximum Gaussian kernel. The data from the six scanning runs were concatenated using the spm_fmri_concatenate function. All analyses reported below were performed in native space on smoothed data.

### MRI data analysis

A separate single-trial GLM was constructed for each participant. Note that group level effects were reported previously by Hill et al. (2020) and are beyond the scope of the current paper. Data from the six study sessions were concatenated and subjected to a ‘least-squares-all’ GLM (Mumford et al., 2014; Rissman et al., 2004) to estimate the BOLD response for each trial separately. Each study event was modeled with a delta function convolved with the canonical hemodynamic response function (HRF). Six regressors representing motion-related variance (three for rigid-body translation and three for rotation) and six session specific regressors were included in each model as covariates of no interest.

For each ROI (see ‘ROI Localization’), we extracted parameter estimates for the single-trial BOLD responses, averaged across all voxels falling within a given ROI. Single-trial BOLD values were used to compute SMEs as the standardized mean difference between subsequently recalled (R) and not recalled (NR) study items using the formula:

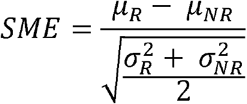

In the above formula, µ_R_ and σ ^2^_R_ refer to, respectively, the across trial mean and variance of BOLD activity for subsequently recalled study items, and µ^2^_NR_ and σ ^2^_NR_ refer to the across trial mean and variance of BOLD activity for subsequently forgotten study items. This formula produces SME values for each ROI that are akin to a Cohen’s *d* effect size estimate. Positive values thus reflect regions where increased brain activity was predictive of subsequent remembering (so-called positive subsequent memory effects) and negative values reflect regions where a relative increase in brain activity is predictive of subsequent forgetting (so-called negative subsequent memory effects).

### iEEG data acquisition and preprocessing

Stereo-EEG data were recorded with a Nihon Kohden EEG-1200 clinical system. Each electrode contained 8-12 contacts spaced 2-4 mm apart. Signals were sampled at 1000 Hz and referenced to a common intracranial contact. Raw signals were subsequently re-referenced to the median white matter signal computed separately for each subject. All analyses were conducted using MATLAB with proprietary and custom-made scripts. We employed kurtosis-based artifact rejection with a threshold of < 5 to exclude interictal activity and abnormal trials (Sederberg et al., 2006). The raw signals were filtered for line noise on a session-by-session basis using a first-order bandstop Butterworth filter with a stopband from 58 to 62 Hz.

### iEEG data analysis

To compute spectral power, we convolved the median white matter re-referenced EEG with 53 complex valued Morlet wavelets (width 6 cycles) spaced logarithmically from 2 to 150Hz. The magnitude of the wavelet transform was then squared and log-transformed to yield instantaneous power. Power estimates for each electrode were z-scored separately for each frequency bin using the mean and standard deviation of the power estimate from the 200 ms pre-stimulus baseline interval. Normalized power was then averaged within six canonical frequency bands: delta (2-4 Hz), theta (4-8 Hz), alpha (8-12 Hz), beta (12-30 Hz), low gamma (30-70 Hz), and high gamma (70-150 Hz). SMEs were computed over the entire 1800 ms epoch during which the study item was presented using the same formula used to compute BOLD SMEs (see above). For subsidiary analyses, additional SMEs were computed separately for early (0-900 ms) and late (900-1800 ms) epochs.

### ROI Localization

Intracranial contacts were localized using post-implant computed tomography (CT) and structural T1 MR scans. CT images were linearly co-registered to the T1 MRI obtained during the fMRI session using FSL FLIRT (FSL version 6.0.1) (Greve & Fischl, 2009; Jenkinson et al., 2002; Jenkinson et al., 2012; Jenkinson & Smith, 2001). For each participant, the native T1 image was then loaded into MRIcron stereotaxic space and overlaid with the co-registered native CT image. As illustrated in Figure 1, microelectrode contacts were visible as high intensity artifacts on the CT overlay. Contacts were manually localized with reference to stereotaxic coordinates in standard MNI space for each patient.

**Figure 1.**
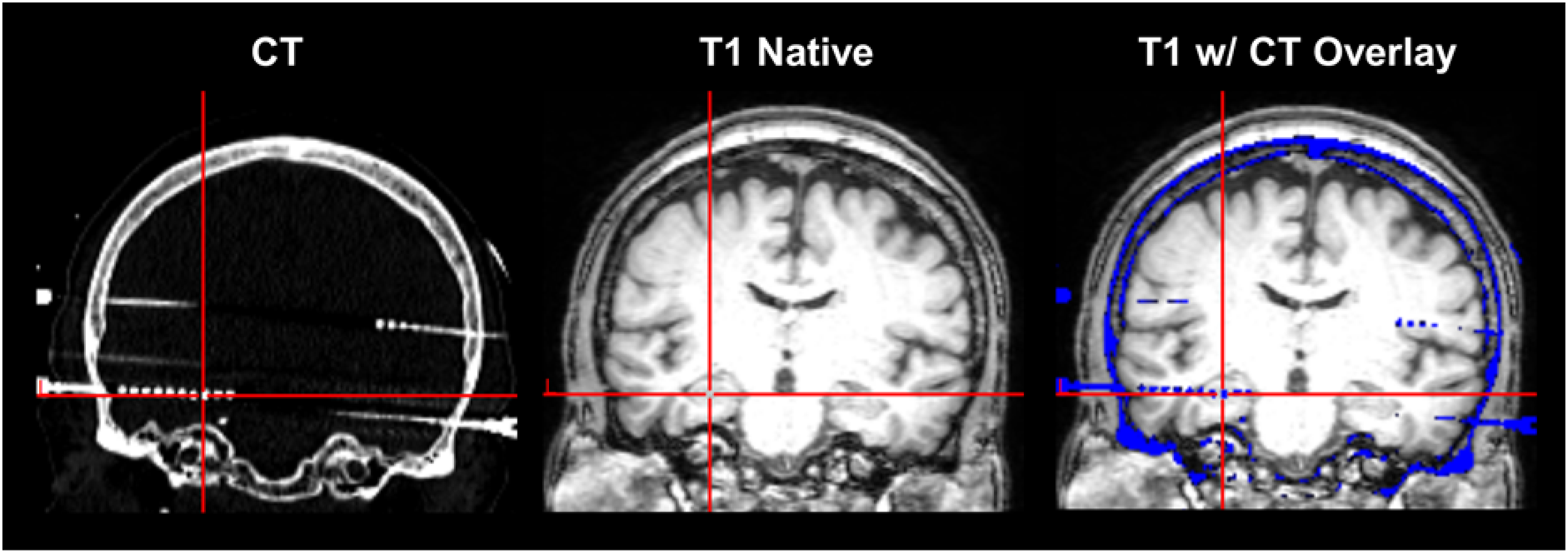
ROI Localization Pipeline. Example of a hippocampal contact localized on a co-registered native CT (left panel) and T1 (middle panel) image. Note that the left and middle panels are for illustrative purposes only. The CT image was overlaid on the T1 image (right panel, CT overlay shown in blue) so that contacts could be manually localized with reference to stereotaxic coordinates in standard MNI space.

Each patient’s native mean functional T2* image was manually inspected to ensure adequate alignment with the native T1 image. To eliminate contacts affected by signal dropout and distortions caused by susceptibility artifacts, we loaded the mean T2* image into MRIcron and visually inspected the coordinates for each contact to ensure adequate signal quality. Contacts falling within areas affected by magnetic susceptibility artifact were flagged and excluded from subsequent analyses. This procedure identified a total of 139 contracts (10%) for exclusion.

To identify contacts located in white matter, tissue segmentation of the structural T1 scans was performed using FAST in FSL (Zhang et al., 2001) with white matter pattern probability set at 70%. Contacts visible on the CT overlay were manually inspected with reference to the white matter mask, and those falling within the mask in all three stereotaxic directions (x, y, z) were labeled as white matter contacts. For each subject, these white matter contacts were combined to provide a grand median reference signal that was used to compute SMEs (see iEEG Data Analysis). We note that the criteria for selecting white matter contacts was more conservative than those for localizing grey matter contacts, ensuring that the white matter reference signal was unlikely to include any residual signal from grey matter. Contacts located outside of the skull were flagged and excluded from further analyses, as were all grey matter contacts showing evidence of ictal activity or other pathology.

For the fMRI analyses, spherical ROIs (3mm radius) were generated using the MarsBaR (v. 0.44) toolbox for SPM. Each ROI was centered on the native stereotaxic coordinates corresponding to the grey matter contacts localized in the aforementioned paragraphs. The mean fMRI BOLD SME was then computed across all voxels falling within each sphere using the procedures described above (see ‘MRI Data Analysis’).

Each contact was labeled by a trained neuroradiologist according to the Automated Anatomical Labelling (AAL) atlas (Tzourio-Mazoyer et al., 2002). For quality assurance, all hippocampal and parahippocampal labeled contacts were also manually inspected and their locations confirmed by the first author. The AAL labels were used to sort ROIs into lobar and sub-lobar parcels in the region-based analyses reported below. The mean number of ROIs for each patient per lobe are reported in Table 1.

**Table 1.** Mean # of ROIs (with range) per subject in each of the four lobar regions.

| Region | Mean (range) # ROIs |
| --- | --- |
| Frontal | 24 (3-55) |
| Temporal | 26 (9-37) |
| Parietal | 14 (2-32) |
| MTL | 11 (4-22) |

### Statistical Analyses

Statistical analyses were carried out using R software (R Core Team, 2017). ANOVAs were conducted using the *afex* package (Singmann et al., 2016) and the Greenhouse-Geisser procedure (Greenhouse & Geisser, 1959) was used to correct degrees of freedom for non-sphericity when necessary. Post-hoc tests on significant effects from the ANOVAs were conducted using the *emmeans* package (Lenth, 2018). Multiple regression and correlation analyses were performed using the *lm* and *cor*.*test* functions in the base R package, respectively. Linear mixed-effects models were performed using the lme4 package (Bates et al., 2015), and degrees of freedom estimated using the Kenward-Roger method. 95% confidence intervals for fixed effects were computed via parametric bootstrapping in the broom.mixed package (Bolker, 2020). All models included a random intercept per subject. Inclusion of additional random intercept and slope terms are described in the relevant sections below. All models were fit using maximum likelihood Laplace approximation, and were refit using restricted maximum likelihood prior to performing nested model comparisons.

## Results

### Behavioral Results

Behavioral results from the fMRI session were previously reported (Hill et al., 2020). The proportion of freely recalled study items from the fMRI session (*M* = .30, *SD* = .11) closely approximated performance during the iEEG session (*M* = .27, *SD* = .09). However, the iEEG session always preceded the fMRI session (*M* = 87 days, *SD* = 66 days). Given the consequent possibility of order effects, and the slight methodological differences between the free recall paradigms administered during the respective sessions (see Methods and Materials), we did not perform a direct statistical test to compare recall performance between the two testing sessions.

### Coupling between BOLD and LFP SMEs varies across brain regions and frequency bands

In the first set of analyses, we examined whether variance in the magnitude of memory-related BOLD signal change could be predicted by variance in memory-related iEEG changes measured from the same anatomical locations, and whether the relationship between BOLD and iEEG effects varied across brain regions. Each ROI was assigned to one of four lobar labels: frontal, temporal, parietal, and medial temporal (including hippocampus, parahippocampal gyrus, and amygdala). Due to sparse coverage, data extracted from ROIs in the occipital lobe (derived from a total of only 8 contacts) were not included in these analyses. For each subject, the across-ROI vector of BOLD SMEs from each of the four lobar regions was entered into the model as the dependent variable. iEEG SMEs recorded from the same ROIs in each frequency band were entered as the fixed effect of interest, along with hemisphere of ictal onset (right, left, bilateral) and handedness (left, right) as nuisance regressors. Using the lobar labels provided for each ROI, region- and subject-wise intercept and slope terms were entered into the respective LME models as fully crossed random effects.

Using nested maximum likelihood ratio tests, we found that, compared to the models with only the subject-level random effects factor, inclusion of the regional random effects significantly improved model fit in each of the six frequency bands (Table 2). These results suggest that the magnitude and/or direction of the relationships between BOLD and iEEG SMEs are regionally variant. Motivated by these findings we specified an additional set of subsidiary LME models separately for each lobar region. Because the number of ROIs per lobe in any given subject was highly variable (Table 1), we elected to perform subject-wise intercept only models (i.e., random intercepts, fixed slopes). The models were otherwise specified as before. Note that modeling the relationship between BOLD and iEEG effects at the level of sub-lobar cortical and subcortical loci (loci here referring to the AAL labels assigned to each ROI) did not explain any additional variance over and above the lobar models (cf. Conner et al., 2011). We therefore report below only the results of the LME models corresponding to each lobar region.

**Table 2.** Comparison of nested random effects.

| Frequency | $X^2$ | $p$-value | ΔAIC |
| --- | --- | --- | --- |
| Delta | 23.73 | $2.84^{-5}$ | 18 |
| Theta | 25.72 | $1.09^{-5}$ | 20 |
| Alpha | 25.34 | $1.31^{-5}$ | 19 |
| Beta | 23.47 | $3.22^{-5}$ | 18 |
| Low Gamma | 53.56 | $1.39^{-11}$ | 48 |
| High Gamma | 29.69 | $1.61^{-6}$ | 24 |

The results of the low and high gamma LME analyses are illustrated in Figure 2. BOLD SMEs positively co-varied with high gamma SMEs in frontal (β = .11, *t* = 2.92, 95% CI = .03, .18), temporal (β = .11, *t* = 3.06, 95% CI = .04, .18), and parietal (β = .26, *t* = 4.86, 95% CI = .15, .36) cortices. BOLD SMEs in the MTL negatively covaried with low gamma SMEs (β = -.14, *t* = -2.39, 95% CI = -.26, -.03). Note that each of these effects remained significant after controlling for the iEEG SMEs in all other frequency bands. Thus, gamma-band power changes explained unique sources of variance in encoding-related BOLD signal change in the neocortex and MTL. BOLD SMEs negatively covaried with theta SMEs in frontal lobe (β = -.12, *t* = -2.35, 95% CI = - .22, -.02) and positively with alpha SMEs in the parietal lobe (β = .21, *t* = 2.98, 95% CI = .07, .34). When controlling for iEEG SMEs in all other frequency bands, only the negative BOLD-theta relationship in the frontal lobe remained significant.

**Figure 2.**
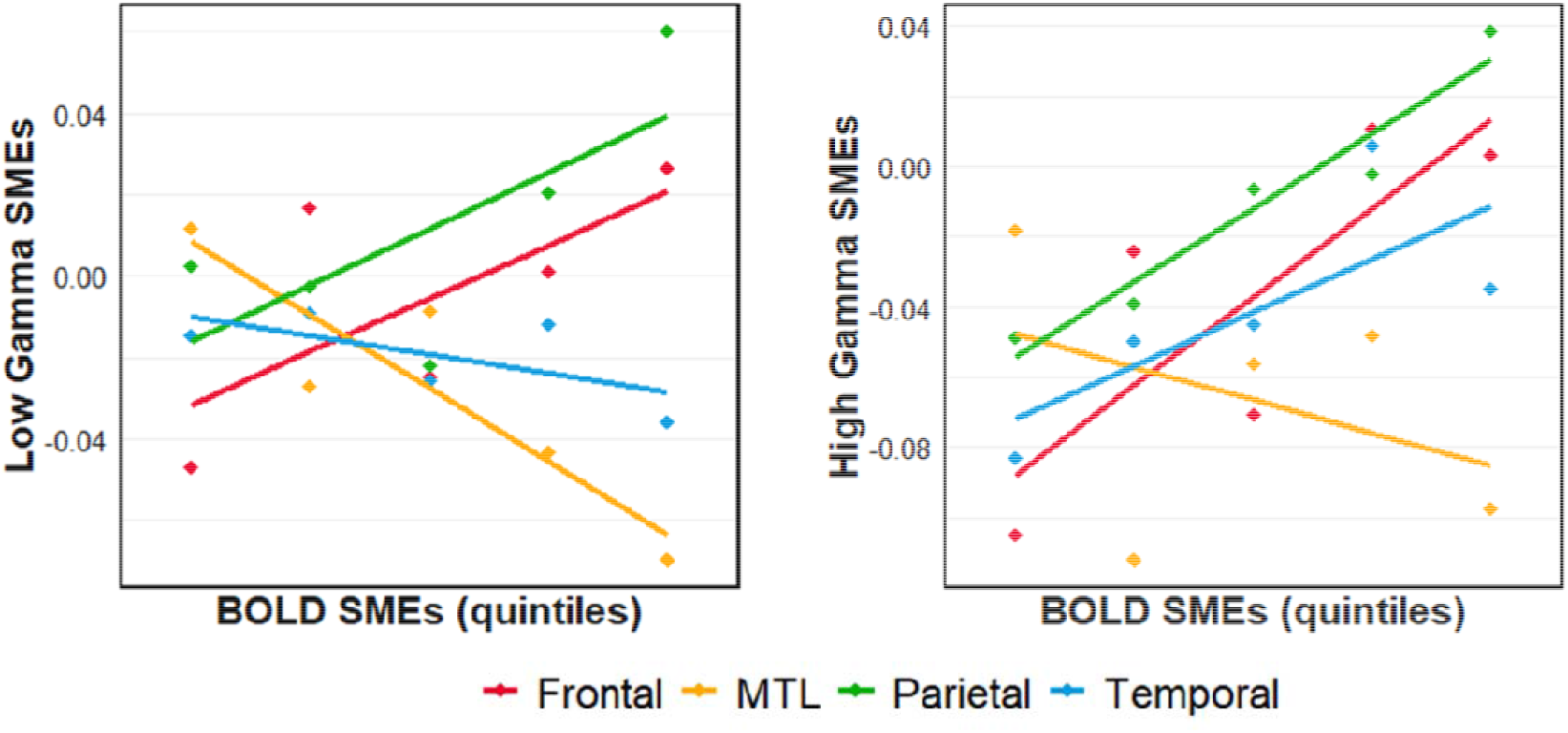
Scatterplots showing the relationship between BOLD and gamma-band subsequent memory effects in the MTL and neocortex. (Left) A significant negative relationship between BOLD and low gamma SMEs was evident in the MTL. The relationship between BOLD and low gamma SMEs in frontal, temporal, and parietal cortex was not significant. (Right) A significant positive relationship between BOLD and high gamma SMEs was evident in frontal, temporal and parietal cortex. These effects were accompanied by a negative but nonsignificant relationship between BOLD and high gamma SMEs in the MTL. Data are binned into quintiles based on the magnitude of BOLD SMEs for visualization purposes.

### Relationship between BOLD and gamma-band SMEs in the hippocampus and parahippocampal gyrus

We next performed a set of subsidiary linear regression analyses to examine whether the relationship between BOLD and iEEG SMEs recorded from the MTL differed between parahippocampal neocortex and hippocampal allocortex (see Introduction). Due to sparse coverage, data extracted from ROIs in the amygdala (derived from a total of only 13 contacts from 5 patients) were not included in these analyses. BOLD SMEs were entered as the dependent variable, and iEEG SMEs, region, and the iEEG x region interaction terms were entered as predictor variables along with hemisphere of ictal onset and handedness as nuisance regressors. The number of ROIs localized to the hippocampus (*M* = 6, range = 0-15) and parahippocampal gyrus (*M* = 5, range = 2-8) was highly variable across subjects. We therefore elected to run linear regression rather than LME analyses, as the error term in the latter can be biased in cases with too few observations per random effect (in this case subject). We note that although these analyses are limited in that ROIs, rather than subjects, are treated as a random effect, a separate set of by-subject LME analyses produced identical results. Thus, for parsimony we report only the results of the linear regression analyses.

The results of the low and high gamma regression analyses are illustrated in Figure 3. The analysis of high gamma effects revealed a significant interaction between region and high gamma SMEs (*F*_(1, 155)_ = 6.21, *p* = .014) which was driven by a negative relationship between BOLD and high gamma SMEs in the hippocampus (*r* = -.22, *p* = .041), and a positive but nonsignificant relationship in the parahippocampal gyrus (*r* = .20, *p* = .132). Regression models for the remaining frequency bands failed to identify any significant region x iEEG interactions (all *p*s > .1). Consistent with the results of the MTL LME analysis reported above, the low gamma model revealed a significant main effect of iEEG (*F*_(1, 159)_ = 11.28, *p* = .001), such that BOLD SMEs negatively covaried with low gamma SMEs recorded from the hippocampus (*r* = - .32, *p* = .002) and tended to do so in the parahippocampal gyrus (*r* = -.21, *p* = .08).

**Figure 3.**
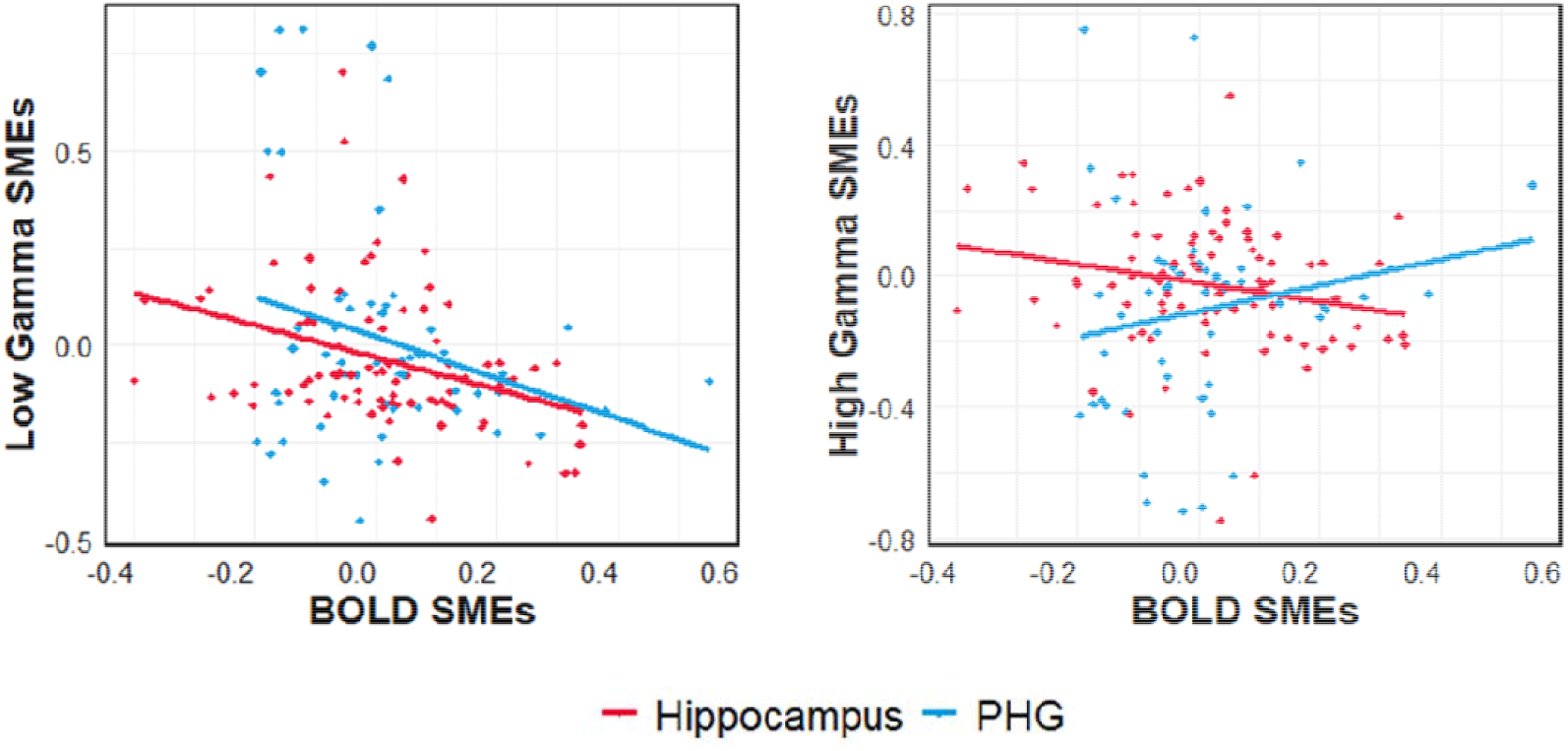
Scatter plots illustrating the relationship between BOLD and gamma-band SMEs in the hippocampus and parahippocampal gyrus. (Left) A significant negative correlation between BOLD and low gamma SMEs was evident in both the hippocampus and parahippocampal gyrus, and the magnitude of these correlations did not differ between the two regions. (Right) A significant negative correlation between BOLD and high gamma SMEs was evident in the hippocampus, accompanied by a positive but nonsignificant correlation between BOLD and high gamma in the parahippocampal gyrus.

We performed a set of follow-up multiple regression analyses with BOLD SMEs as the dependent variable, and the relevant gamma-band iEEG SME (low, high), ROI hemisphere, and the iEEG x ROI hemisphere interaction term as predictor variables, along with handedness and hemisphere of ictal onset as nuisance regressors. The analysis of gamma-band effects in the hippocampus revealed nonsignificant interactions between hemisphere and low gamma (*F*_(1, 86)_ = 0.63, *p* = .429) and high gamma (*F*_(1, 86)_ = 0.00, *p* = .968) SMEs. In the parahippocampal gyrus, there was a significant interaction between hemisphere and high gamma SMEs (*F*_(1, 58)_ = 7.10, *p* = .010) which was driven by a robust positive BOLD-iEEG relationship in the left hemisphere (*r* = .52, *p* = .002) accompanied by a nonsignificant relationship in the right hemisphere (*r* = -.14, *p* = .477). The interaction between hemisphere and low gamma SMEs in the parahippocampal gyrus was not significant (*F*_(1, 58)_ = 0.11, *p* = .740).

### Frontal BOLD effects are differentially predicted by early and late components of delta- and theta-band activity

In the foregoing analyses, iEEG SMEs were computed over the entire 1800 ms encoding period during which a study word was displayed. Although this roughly approximated the sampling rate of fMRI volume acquisition (2000 ms), it risks collapsing across meaningful temporal variation in the electrophysiological effects. Therefore, in a final set of analyses, we examined whether the relationship between BOLD and iEEG effects differed when iEEG SMEs were estimated for early (0-900) and late (900-1800) encoding epochs. We specified LME models separately for each lobar location using an approach similar to that described in previous sections. For each subject, the across-ROI vector of BOLD SMEs from a given lobar region was entered into the model as the dependent variable. Early and late iEEG effects, epoch (early, late), and the iEEG x epoch interaction were entered as fixed effects of interest, along with handedness and hemisphere of ictal onset as nuisance regressors. Subject-wise intercepts were entered as a random effect (i.e., random intercepts, fixed slopes). Given our a priori interest in hippocampal effects, we also performed linear regression analyses on hippocampal BOLD and iEEG SMEs separately for each frequency band. For the multiple regression analyses, BOLD SMEs were entered into each respective model as the dependent variable, and iEEG SMEs, epoch, and the iEEG x epoch interaction term were entered as predictor variables along with handedness and hemisphere of ictal onset as covariates of no interest.

Modelling the relationship between BOLD and iEEG effects in the frontal cortex revealed significant interactions between epoch and low-frequency SMEs in both the delta (*F*_(1, 695.79)_ = 8.60, *p* = .003) and theta (*F*_(1, 694.59)_ = 6.32, *p* = .012) frequency bands (Figure 4). Post-hoc analyses of the delta-band effects revealed a significant *positive* relationship between BOLD and delta SMEs during the late epoch (β = .19, *t* = 3.84, 95% CI = .09, .29), along with a negative but nonsignificant relationship during the early epoch (β = -.06, *t* = -1.32, 95% CI = -.15, .03). By contrast, post-hoc analyses of the theta-band effects revealed a significant *negative* relationship between BOLD and theta SMEs during the early epoch (β = -.18, *t* = -3.67, 95% CI = -.27, -.08), along with a positive but nonsignificant relationship during the late epoch (β = .07, *t* = 1.21, 95% CI = -.04, .17). The early and late temporal epochs did not moderate the relationship between BOLD and iEEG effects in any of the remaining lobar models (all *p*s > .08). Nor did we observe any evidence that epoch moderated the relationship between BOLD and iEEG effects in the hippocampus (all *p*s > .4).

**Figure 4.**
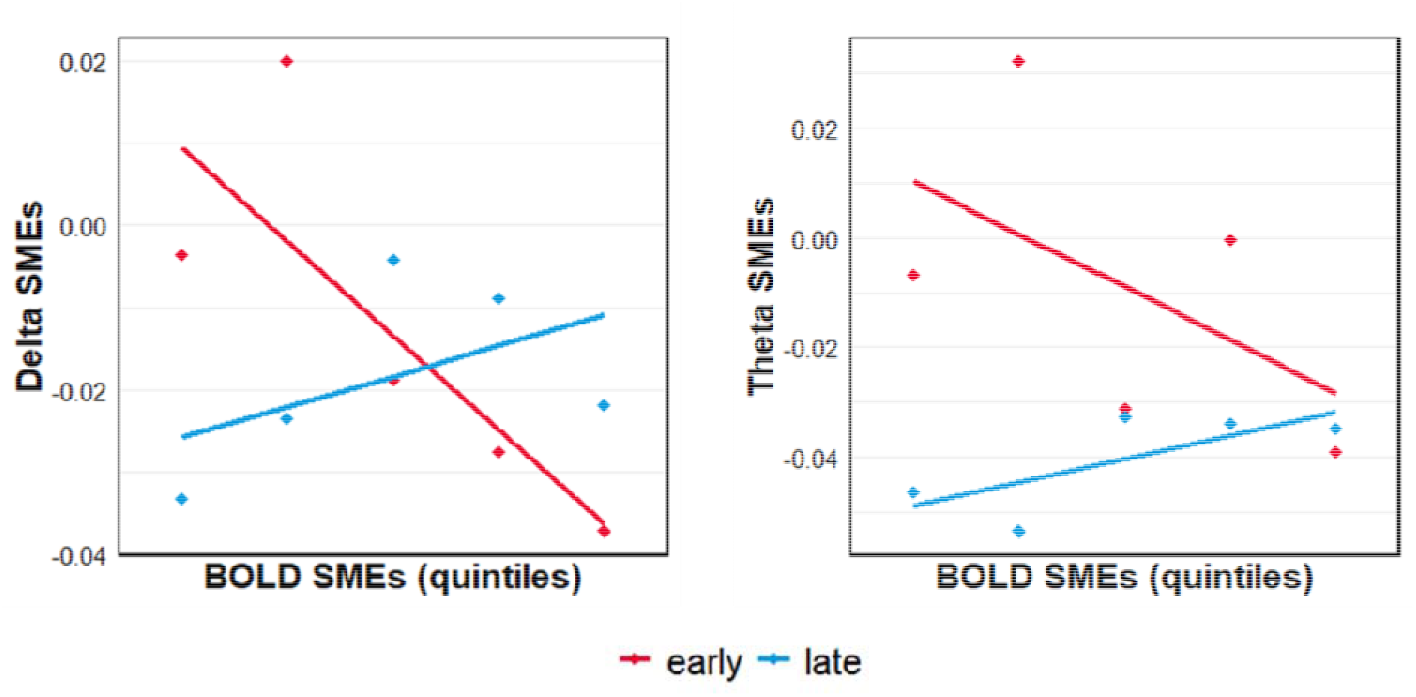
Scatter plots showing the relationship between BOLD and low frequency iEEG SMEs that were moderated by epoch (early, late) in the frontal cortex. Data are binned into quintiles based on the magnitude of BOLD SMEs for visualization purposes.

## Discussion

We examined whether encoding-related differences in electrophysiological activity could predict analogous differences in fMRI BOLD signal magnitude, and whether any such relationships between these neurophysiological and hemodynamic signals varied according to region. BOLD and high gamma (70-150 Hz) SMEs positively covaried across much of the neocortex, with reliable relationships evident in frontal, temporal, and parietal cortices. Notably, this relationship was reversed in the hippocampus, where a negative correlation between BOLD and high gamma SMEs was evident. We also observed a negative relationship between BOLD and low gamma (30-70 Hz) SMEs in the MTL more broadly. In frontal cortex, low-frequency delta (2-4 Hz) and theta (4-8 Hz) activity explained unique variance in BOLD SMEs, and these effects were moderated by epoch (early vs. late). Below, we discuss the significance of these findings in respect of regional variability in the transfer function between neural activity and the fMRI BOLD signal.

As just noted, using the subsequent memory procedure (Paller & Wagner, 2002), we identified robust coupling between encoding-related modulation of high gamma power and BOLD signal amplitude across much of the neocortex, including frontal, temporal, and parietal cortices. The relationship between BOLD and high gamma SMEs did not vary at the level of sub-lobar cortical loci. These findings are notable for two reasons. First, the regionally invariant relationship between BOLD and high gamma effects across much of the neocortex observed in the present study is consistent with numerous prior reports of preferential coupling between BOLD and high frequency iEEG activity measured from primary sensory, motor, and association cortex in behaving humans (Conner et al., 2011; Hermes et al., 2012; Nir et al., 2007; Ojemann et al., 2010). Second, the present findings replicate and extend these prior studies by establishing a link between modulation of BOLD and high frequency iEEG activity during a memory encoding task.

In stark contrast to the robust positive relationships observed across much of the neocortex, we identified a negative relationship between BOLD and both low and high gamma SMEs in the hippocampus. Moreover, the negative relationship between BOLD and high gamma SMEs observed in the hippocampus was dissociable from the relationship evident in anatomically proximal MTL neocortex. These findings are consistent with the proposal that regional variability in patterns of coupling between BOLD and high gamma SMEs reflect regional differences in neurovascular coupling, specifically, between the hippocampus and neocortex (for review, see Ekstrom, 2021). Sparse coding and vascularization schemes might explain the existence of a *null* relationship between BOLD and gamma-band iEEGs in the hippocampus relative to the neocortex (should that have been observed), but such factors cannot readily account for the reliable *negative* relationships that were actually observed for both low and high gamma effects in the present study. Sparse firing of principal cells in the hippocampus (particularly in dentate gyrus and CA3) is made possible by dense recurrent inhibitory interneurons that promote pattern separation (Yassa & Stark, 2011). Because inhibition is metabolically costly, it may be that it is these signals that were responsible for heightened hippocampal BOLD responses, while simultaneously down-regulating high frequency iEEG signals. This account might also explain why variation in the firing of sparsely distributed principal neurons in the hippocampus can seemingly be associated with the robust hippocampal BOLD effects that are evident across a variety of behavioral tasks such as memory encoding (Kim, 2011; Spaniol et al., 2009) and spatial navigation (e.g., Doeller et al., 2008). Alternatively, sparse capillary density in the hippocampus (Borowsky & Collins, 1989) might produce situations in which stimulus evoked increases in brain activity and oxygen consumption (CMRO_2_) outpace the regional supply of oxygenated hemoglobin, leading to a negative BOLD response reflective of increased venous deoxygenated hemoglobin despite an increase in neural activity (Schriddle et al., 2008; see also Ances et al., 2008).

BOLD SMEs in the hippocampus negatively covaried with both low and high gamma SMEs recorded from the same locations. Low gamma effects remained significant when controlling for concurrent high gamma SMEs (though the high gamma effect was rendered nonsignificant when controlling for concurrent low gamma effects). This functional dissociation between negative BOLD effects and low and high gamma is consistent with prior research reporting that low and high gamma LFPs are distinct in both their neurophysiological correlates (Buzsaki & Wang, 2012; Colgin et al., 2009; Ray & Maunsell, 2011) and their functional significance (Bieri et al., 2014; Colgin, 2015; Colgin & Moser, 2010). The present findings thus extend much of the rodent work to humans while providing novel evidence for unique low and high gamma components to the hippocampal BOLD signal. We remain agnostic, however, as to the neurophysiological significance of these effects, and acknowledge that future work is needed to elucidate whether low and high gamma effects do indeed reflect distinct neural correlates of the hippocampal BOLD signal.

In the frontal cortex, BOLD SMEs were related to low frequency delta and theta SMEs and, for each frequency band, this relationship was moderated by encoding epoch (early vs. late). As is illustrated in Figure 4, both the delta- and theta-band effects were characterized by a negative relationship with BOLD during the early epoch, accompanied by a modest positive relationship during the later epoch (though the reliability of these effects differed as a function of frequency band and epoch). We caution that because these results were unanticipated, they should be interpreted cautiously and are clearly in need of replication.

Due to safety considerations, simultaneous iEEG and fMRI recordings are not readily obtainable in humans. We therefore obtained electrophysiological and hemodynamic recordings from the same individuals in sequential experimental sessions, raising the possibility that order or practice effects may have confounded behavioral performance during the fMRI session. Another potential limitation of the present study concerns the methodological differences between the free recall paradigms employed during the fMRI and iEEG sessions. Study lists in the fMRI session comprised 15 concrete nouns compared to the 10 or 12 study items employed in the iEEG sessions. The length of the distractor interval also differed between the fMRI (15s) and iEEG (20s) sessions, as did the amount of time each study item was presented on the screen (1800 vs. 1600 ms for the fMRI and iEEG sessions, respectively). Variability in each of these task parameters has been shown to influence free recall performance (Murdock et al., 1962; Roberts et al., 1972; Ward, 2002). Although we are encouraged by the similar behavioral performance observed during the fMRI and iEEG versions of the task, we are unable to definitively rule out the possibility that these task discrepancies impacted the relationship between the two classes of SME.

Experimental applications of iEEG are currently limited to patients with medically refractory epilepsy, introducing potential constraints on the generalizability of intra-cerebral findings. Leveraging the noninvasiveness afforded by fMRI, we recently reported that group level BOLD SMEs in the same TLE patient cohort reported here did not reliably differ from SMEs observed in an age-matched neurologically healthy control group (Hill et al., 2020). Thus, neuropathology associated with TLE was apparently insufficient to give rise to detectable differences in the functional neuroanatomy of episodic memory encoding as this is reflected by the fMRI BOLD signal. These findings do not, however, rule out the possibility that coupling between electrophysiological and BOLD effects might be altered by disease status. We note that this issue cannot be resolved using within subjects designs owing to the aforementioned invasiveness of iEEG.

In conclusion, we identified a robust positive relationship between encoding-related BOLD and high gamma activity in frontal, temporal, and parietal cortex, replicating findings from numerous prior studies (for reviews, see Ekstrom, 2021; Ojemann et al., 2013). Importantly, this relationship was reversed in the hippocampus, where BOLD SMEs negatively covaried with both low and high gamma SMEs. These results suggest that the neurophysiological bases of the BOLD signal in the hippocampus differ from those in the neocortex.

## Notes

**Funding**: This project was supported by the National Institute of Neurological Disorders and Stroke (grant number R21NS095094)

### Competing Interest Statement

The authors have declared no competing interest.

